# SKN-1 is a metabolic surveillance factor that monitors amino acid catabolism to control stress resistance

**DOI:** 10.1101/2022.02.03.479044

**Authors:** Phillip A. Frankino, Talha F. Siddiqi, Theodore Bolas, Raz Bar-Ziv, Holly K. Gildea, Hanlin Zhang, Ryo Higuchi-Sanabria, Andrew Dillin

## Abstract

The deleterious potential to generate oxidative stress and damage is a fundamental challenge to metabolism. The oxidative stress response transcription factor, SKN-1/NRF2, can sense and respond to changes in metabolic state, although the mechanism and physiological consequences of this remain unknown. To explore this connection, we performed a genetic screen in *C. elegans* targeting amino acid catabolism and identified multiple metabolic pathways as regulators of SKN-1 activity. We found that genetic perturbation of the conserved amidohydrolase *T12A2.1/amdh-1* activates a unique subset of SKN-1 regulated detoxification genes. Interestingly, this transcriptional program is independent of canonical P38-MAPK signaling components but requires the GATA transcription factor ELT-3, nuclear hormone receptor NHR-49, and mediator complex subunit MDT-15. This activation of SKN-1 is dependent on upstream histidine catabolism genes HALY-1 and Y51H4A.7/UROC-1 and may occur through accumulation of a catabolite, 4-imidazolone-5-propanoate (IP). Triggering SKN-1 activation results in a physiological trade off of increased oxidative stress resistance but decreased survival to heat stress. Together, our data suggest that SKN-1 is a key surveillance factor which senses and responds to metabolic perturbations to influence physiology and stress resistance.

## INTRODUCTION

Metabolism is central to normal cell function and is dysregulated in human diseases such as metabolic syndrome, diabetes, and cancer (DeBerardinis and Thompson, 2012). In the most basic sense, metabolism is the sum of all biochemical reactions in the cell, including reactions that create or break down complex molecules (anabolism and catabolism, respectively). The catabolism of amino acids leads to the accumulation of breakdown products, or catabolites, that are essential for creating cellular energy through the tricarboxylic acid (TCA) cycle and cellular respiration (Martínez-Reyes and Chandel, 2020). Outside of their role in creating cellular energy, many of these catabolites have been identified as signaling molecules that affect both normal cell function and disease. For example, tryptophan catabolites are known immunomodulators, and elevated expression of tryptophan catabolism enzymes is associated with cancer progression and poor prognosis (McGaha et al., 2012). Additionally, the histidine catabolite, imidazolone propionate, is elevated in type 2 diabetic patients and has been shown to impair insulin signaling (Chong et al., 2018; Molinaro et al., 2020). Despite their importance, our understanding of the identity, mechanism and physiological consequences of catabolite signaling is incomplete.

A key challenge for metabolism is the resolution of deleterious byproducts that can damage cellular components. For example, the electron transport chain of the mitochondria is the main site of ATP generation but also produces harmful reactive oxygen species (ROS), a form of oxidative stress that damages DNA, lipid membranes and proteins (Nolfi-Donegan et al., 2020). To resolve oxidative damage, cells have evolved a conserved stress response controlled by the transcription factor SKN-1/NRF2 that upregulates antioxidant synthesis and detoxification enzymes to neutralize oxidants and export toxic molecules from the cell (Blackwell et al., 2015). In the nematode *C. elegans*, the oxidative stress response (OxSR) is initiated in response to oxidative damage through a signaling cascade that converges on the conserved map kinase (MAPK) pathway, resulting in the phosphorylation and activation of the MAPKK and P38/MAPK homologs (SEK-1 and PMK-1, respectively) (Inoue et al., 2005). Once this signaling cascade is initiated, nuclear factors such as ELT-3, NHR-49, and MDT-15 are required for upregulation of stress response targets (Goh et al., 2014, 2018; Hu et al., 2017; Wu et al., 2016). Together, the OxSR alleviates oxidative stress and restores homeostasis to promote cell and organismal survival.

Historically, SKN-1 is known as the master regulator of the OxSR but emerging literature has implicated it as a metabolic surveillance factor. For example, exogenous supplementation of amino acids are sensed and activate SKN-1-mediated transcription (Edwards et al., 2015). SKN-1 can also respond to changes in proline catabolism to mobilize lipids during starvation (Pang et al., 2014). Furthermore, genetic perturbation of multiple amino acid catabolic pathways activates SKN-1-mediated transcription (Fisher et al., 2008; Ravichandran et al., 2018; Tang and Pang, 2016). Intriguingly, these instances of SKN-1 activation may involve diverse catabolites, integrating the state of multiple metabolic pathways to sense and respond to metabolic imbalance through a single effector. To date, no study has comprehensively probed metabolic pathways to understand the surveillance and response role of SKN-1 in metabolism.

Here, we identify multiple pathways of amino acid catabolism that, when perturbed, activate a distinct transcriptional response driven by SKN-1. Using a mutant of histidine catabolism as a model, we show that this response is independent of canonical MAPK signaling pathways and may partially depend on GCN2 and mTOR homologs *gcn-2* and *let-363*. We also demonstrate the necessity of nuclear factors previously implicated in the OxSR for SKN-1 activation. Interestingly, this response is dependent on the upstream enzymes of the histidine catabolism pathway, suggesting endogenous catabolites may activate SKN-1. Activation of SKN-1 via mutation of the histidine catabolism pathways results in increased oxidative stress resistance at the cost of decreased resistance to heat stress indicating that SKN-1 mediates a tradeoff between stress response survival. Together, our data uncover a novel metabolic surveillance mechanism driven by SKN-1, which likely works through accumulation of catabolite intermediates, to control susceptibility to stress.

## RESULTS

### Genetic perturbation of amino acid catabolism pathways activates SKN-1

To uncover the genetic mechanisms by which SKN-1 serves as a metabolic surveillance factor, we performed an RNAi screen to comprehensively survey amino acid catabolism pathways that, when perturbed, activate a SKN-1 dependent transcriptional response. We constructed a sub-library containing 78 RNAi clones targeting catabolic pathways for all of the 20 proteinogenic amino acids including genes involved in glutathione (GSH) synthesis (table 1). Using this RNAi sub-library, we assessed SKN-1 activity by measuring the fluorescence of animals expressing GFP downstream of the *gst-4* promoter (*gst-4p::GFP*), a well established reporter of SKN-1 (Link and Johnson, 2002). Notably, we found that knockdown of glutathione synthesis, tyrosine or valine catabolism enzymes activate the SKN-1 reporter as previously described, confirming the ability of our screen to identify known regulating enzymes of, and pathways surveilled by, SKN-1 (fig. 1a, table 1) (Fisher et al., 2008; Wang et al., 2010). Interestingly, we found that genetic perturbation of histidine, glycine and phenylalanine catabolism led to activation of SKN-1 (fig. 1a, table 1). Together, these results suggest that SKN-1 is a metabolic regulator which responds to changes in multiple amino acid catabolism pathways, and possibly directly to amino acid levels.

**Figure 1.**
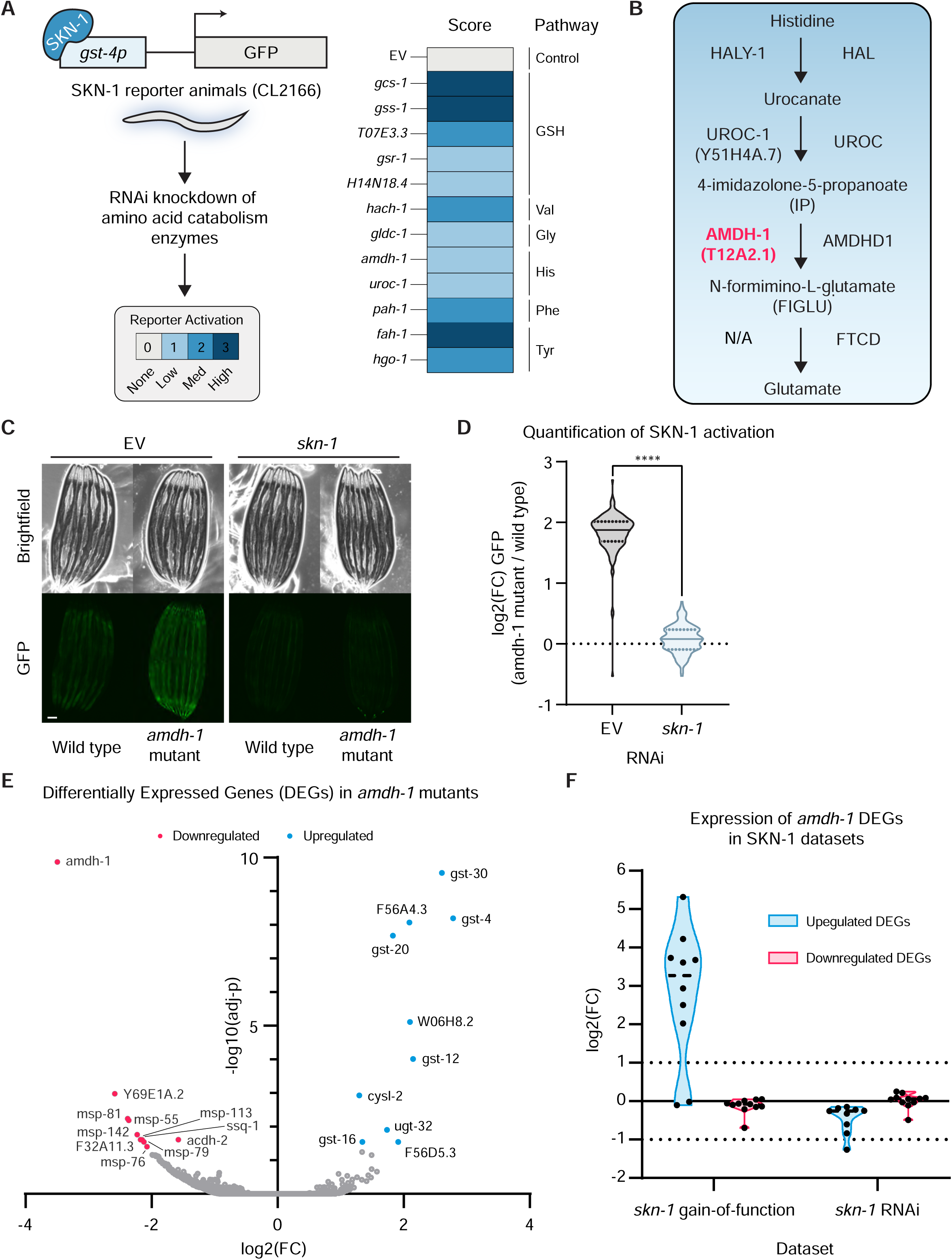
Perturbation of histidine catabolism activates a SKN-1 mediated detoxification response. (A) Experimental scheme of RNAi screen to uncover amino acid catabolism enzymes that affect SKN-1 activation (left), and scores of tested genes (right). RNAi knockdown of genes that suppressed the reporter were scored but not included (table 1). (B) Enzymes and intermediates involved in the histidine catabolism pathway, in *C. elegans* (left) and humans (right) (“N/A” represents no identified *C. elegans* enzyme for this step). (C) Fluorescent images of SKN-1 transcriptional reporter, *gst-4p::GFP*, in wildtype or mutant backgrounds on RNAi Scale bar, 100 μm (D) Quantification of SKN-1 activation (*amdh-1* mutant normalized to median of wild type) from (C), Data are representative of n = 3 biological replicates, n > 121 animals per replicate, **** = P < 0.0001 using a Mann-Whitney two-tailed test. (E) Volcano plot of genes in *amdh-1(uth29)* compared to N2 wildtype control. Differentially expressed genes (DEGs) shown in red (downregulated) and blue (upregulated), adjusted-p < 0.05 (F) Meta analysis of DEGS from *amdh-1(uth29)* mutants in gain of function *skn-1(lax188)* and *skn-1* RNAi datasets (Nhan et al., 2019; Steinbaugh et al.).

### Knockout of histidine catabolism enzyme AMDH-1 triggers a SKN-1-mediated detoxification response

Our screen revealed that the phenylalanine, glycine and histidine catabolism pathways are surveilled by SKN-1, and activate this transcription factor when perturbed. Both phenylalanine, through the phenylalanine hydroxylase *pah-1*, and glycine catabolism, through the glycine cleavage protein *gldc-1*, have established connections to SKN-1 via the NADPH oxidase *bli-3* and antioxidant synthesis, respectively (Calvo et al., 2008). We chose the conserved histidine catabolism pathway as a model to further elucidate this mechanism of SKN-1 activation, as histidine is an essential amino acid and its metabolic pathway remains largely uncharacterized in *C. elegans. T12A2.1*, which was renamed **<u>AM</u>**<u>i</u>**<u>D</u>**o**<u>H</u>**ydrolase domain containing protein **<u>1</u>** (*amdh-1)*, is a conserved amidohydrolase in the histidine catabolism pathway that processes its substrate, 4-imidazolone-5-propanoate (IP), to create N-formimino-L-glutamate (FIGLU) (fig 1b). To further explore the mechanism of SKN-1 surveillance of amino acid catabolic processes, we used CRISPR-Cas9 to generate a putative null allele, *amdh-1*(*uth29)*, by introducing a premature stop codon in exon 1 of its coding sequence (supplemental figure 1). We found that *amdh-1* mutants robustly induce the SKN-1 reporter strain in a SKN-1-dependent manner, compared to wild-type animals (fig 1c,d). To determine whether *amdh-1* mutation altered global SKN-1 transcription, we analyzed gene expression changes in *amdh-1* mutants versus wild-type animals (fig 1e, table S1). We found that a unique subset of SKN-1 targets, the detoxification enzyme family of glutathione-s-transferases (GSTs), were among the most upregulated genes in our dataset, suggesting that changing AMDH-1 levels triggers a specific transcriptional output, likely driven by SKN-1. Direct comparison of differentially expressed genes (DEGs) in our dataset to previously published datasets revealed that genes upregulated in *amdh-1* mutants were also highly upregulated in *skn-1(lax188)* gain-of-function animals and downregulated in worms treated with *skn-1* RNAi (fig 1f) (Nhan et al., 2019; Steinbaugh et al.). In contrast, downregulated genes in *amdh-1* mutant animals were not differentially regulated under *skn-1* loss or gain-of-function, suggesting that AMDH-1 may also impact SKN-1-independent processes. Interestingly, knockdown of *amdh-1* did not affect the expression of another well-characterized SKN-1 reporter (*gcs-1p::GFP*), further suggesting that there are distinct transcriptional responses modulated by SKN-1 (supplemental fig 1b,c). Taken together, our data show that perturbation of histidine catabolism upregulates a specific transcriptional program driven by SKN-1, that is only a subset of the general oxidative stress response, and is likely tailored for the perturbation of histidine catabolism. This response includes detoxification enzymes but not other known SKN-1 targets such as antioxidant synthesis enzymes.

### SKN-1 activation in *amdh-1* mutants is dependent on known oxidative stress regulators but not canonical p38/MAPK or nutrient signaling pathways

Amino acids play essential roles in maintaining energy homeostasis, a process controlled by a few key nutrient regulators (Efeyan et al., 2015). Therefore, we hypothesized that perturbation of histidine catabolism, and thus SKN-1 activation, may be tightly regulated by these nutrient sensing pathways. To test this hypothesis, we knocked down orthologs of nutrient regulators mTORC1/2 (*let-363)*, RRAGA *(raga-1*), AMPK (*aak-2)*, FOXO (*daf-16*), TFEB (*hlh-30*) and GCN2 (*gcn-2*) and surveyed SKN-1 activation in *amdh-1* mutants. Strikingly, we observed that none of these regulators were completely required for SKN-1 activation (supplemental figure 2a,b). Knockdown of *aak-2* had no effect while *daf-16, hlh-30* or *raga-1* further increased SKN-1 activation in *amdh-1* mutants. Notably, *gcn-2* or *let-363* knockdown significantly decreased SKN-1 activation in these mutants, although SKN-1 levels remained over 2 fold greater (log2(FC) > 1) in *amdh-1* mutants compared to wild-type animals fed RNAi against these genes (supplemental figure 2a,b). Interestingly, we observed increased activation of SKN-1 in *raga-1* knockdown conditions, targeting RRAGA/mTORC1, and suppression of activation upon *let-363* knockdown, targeting both mTORC1/mTORC2. These data suggest that activation of SKN-1 in *amdh-1* mutants may depend on GCN2 and mTORC2 while it is independent of oter nutrient regulators AMPK, FOXO, TFEB and mTORC1/RRAGA.

In *C. elegans*, SKN-1 can be activated via phosphorylation by the p38/MAPK ortholog, PMK-1, in a signaling cascade that requires the MAPKK, SEK-1 (Inoue et al., 2005). To assess the requirements of these well established regulators on SKN-1 activation, we tested whether animals with mutations in this pathway can still activate SKN-1 upon *amdh-1* knockdown. We observed that *pmk-1(km25)* and *sek-1(km4)* mutants fail to suppress SKN-1 activation and instead exhibit increased activation with *amdh-1* knockdown (fig 2a,b). These data show that canonical MAPK regulators are not required for SKN-1 activation in the face of metabolic perturbations and may even negatively regulate this response.

**Figure 2.**
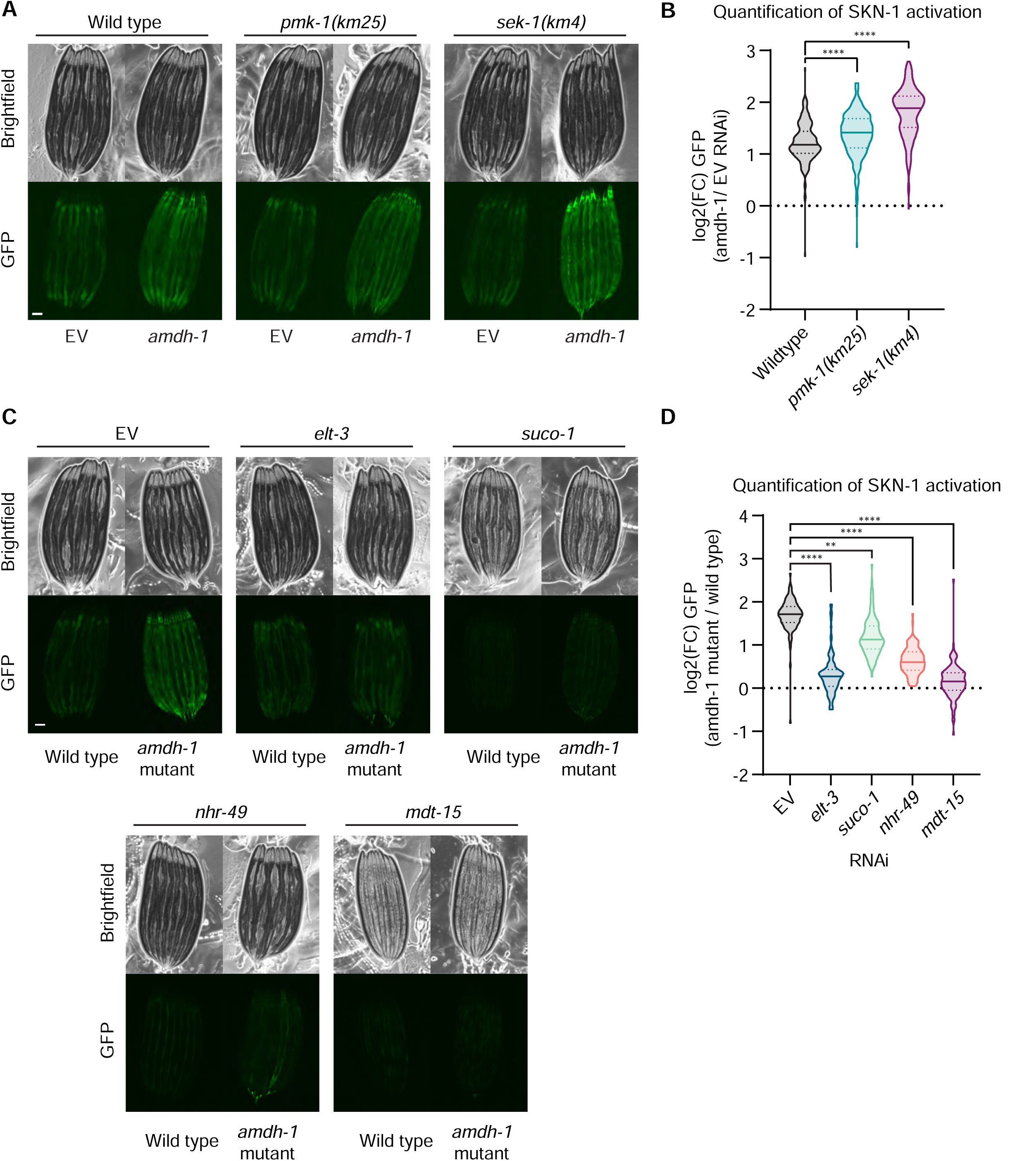
Genetic requirements of SKN-1 activation upon perturbation of histidine catabolism. (A) Fluorescent images of SKN-1 reporter animals in a wildtype, *sek-1(km4)* or *pmk-1(km25)* mutant animals fed *amdh-1* RNAi. Scale bar, 100 μm. (B) Quantification of SKN-1 activation (*amdh-1* RNAi normalized to median of EV) from (A), Data shown are representative of n = 3 biological replicates with n > 172 animals per condition for each replicate. **** = P < 0.0001 using a one-way ANOVA. (C) Fluorescent images of SKN-1 reporter animals in a wildtype or *amdh-1(uth29)* mutant background fed RNAi targeting *elt-3, suco-1, nhr-49*, and *mdt-15*. Scale bar, 100 μm. (D) Quantification of SKN-1 activation (*amdh-1* mutant normalized to median of wild type) from (C), Data shown are representative of n = 3 biological replicates with n > 88 animals per condition for each replicate. ** = P < 0.01, **** = P < 0.0001 using a one-way ANOVA.

To identify genetic pathways required for SKN-1 activation in *amdh-1* mutants, we performed a genetic suppressor screen using EMS mutagenesis on SKN-1 reporter animals in an *amdh-1(uth29)* background. Using a combination of backcrossing and deep sequencing as previously described (Lehrbach et al., 2017), we identified 6 alleles in 4 genes that are required for SKN-1 activation in *amdh-1* mutants (supplemental figure 2c). Among the mutants identified was a putative DNA binding domain mutant of *skn-1* that is present across all four isoforms (supplemental figure 2c). These worms are slow growing, likely due to the requirement for SKN-1 in development (Bowerman et al., 1992). The discovery of this new SKN-1 allele validates the screen and supports the previous finding that this phenotype is dependent on *skn-1* (figure 1c,d). Among the remaining mutations found in our screen, we identified a novel SKN-1 regulator, *suco-1*, and a previously identified regulator, *elt-3*, as suppressors of SKN-1 activity in *amdh-1(uth29)* mutant animals (supplemental figure 2c). RNAi knockdown of *suco-1* or *elt-3* suppress SKN-1 activation in *amdh-1* mutants, phenocopying the EMS mutants and suggesting a causal relationship between these regulators and SKN-1 (fig2c,d). *suco-1* encodes an ortholog of the SLP/EMP65 complex identified in yeast to function in ER protein homeostasis, a process previously implicated in the OxSR (Glover-Cutter et al., 2013; Zhang et al., 2017). *elt-3* is a required factor for induction of OxSR gene expression (Hu et al., 2017). The existing role of *elt-3* in the OxSR prompted further investigation of other required nuclear factors.

SKN-1 drives expression of detoxification enzymes, such as GST-4, under conditions of oxidative stress with the help of nuclear factors like ELT-3, nuclear hormone receptor NHR-49, and mediator complex subunit, MDT-15 (Goh et al., 2014, 2018; Hu et al., 2017; Wu et al., 2016). As these nuclear factors are well established to be required for both stress response gene transcription and survival upon oxidative stress, we hypothesized that they may also be required for SKN-1 activation in *amdh-1* mutants. We found that knockdown of *nhr-49* or *mdt-15* significantly suppressed the activation of SKN-1 in *amdh-1(uth29)* mutants (fig 2c,d). Together, these data suggest that activation of SKN-1 in animals with perturbed histidine catabolism is dependent on canonical nuclear regulators of the OxSR.

### Enzymes HALY-1 and Y51H4A.7/UROC-1 are required for SKN-1 activation in AMDH-1 mutants

Through the EMS suppressor screen, we also identified 3 independent alleles of *haly-1 (uth92, uth93, uth95)*, the conserved histidine ammonia lyase that is rate limiting for the histidine catabolism pathway, as suppressors of SKN-1 activity (fig 3a, supplemental figure 2c). Expression of a wild-type copy of *haly-1* rescued the suppression of SKN-1 activation in *haly-1(uth92)* and *haly-1(uth93)* mutants, confirming the causative nature of these mutations (supplemental figure 3a,b).

**Figure 3.**
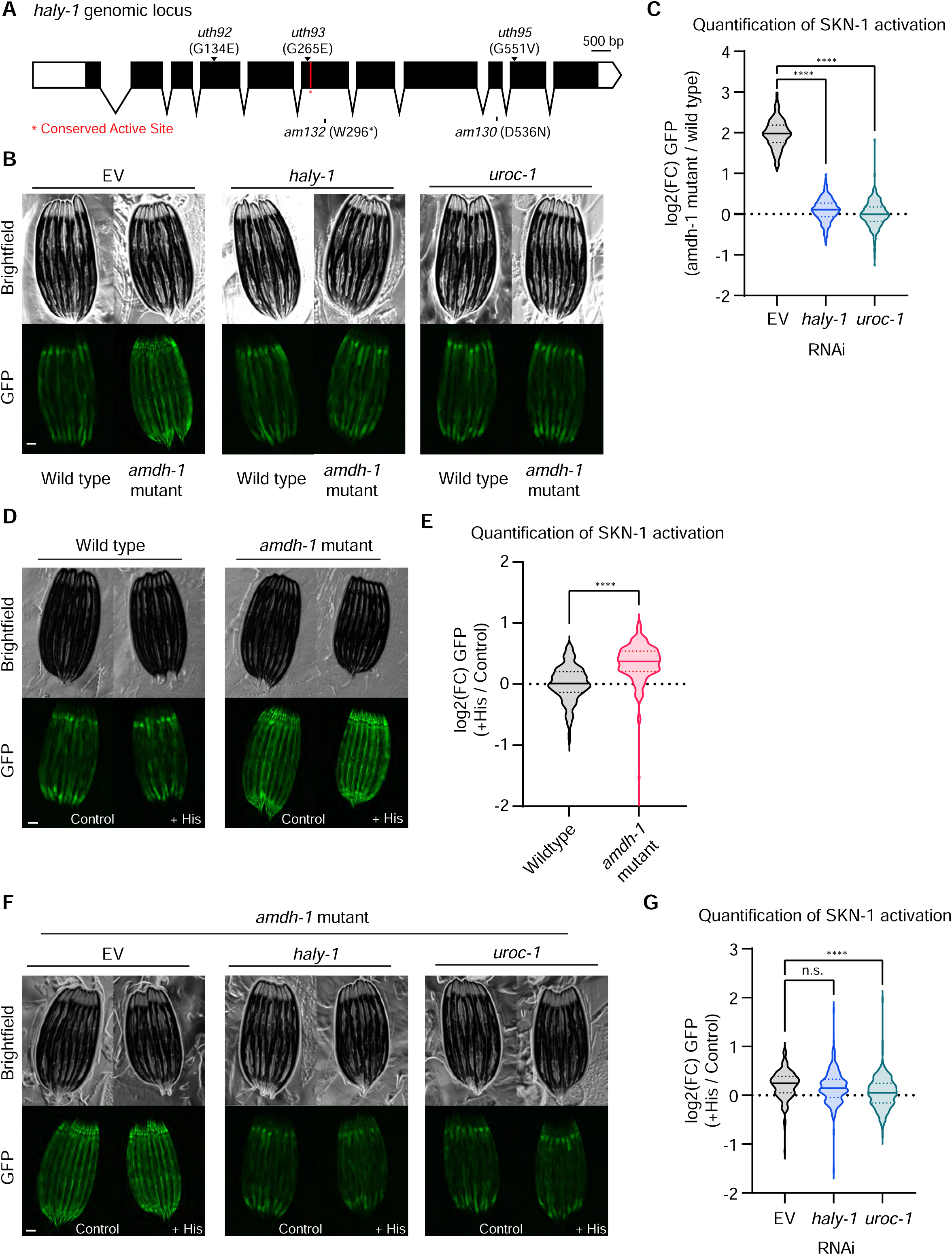
Activation of SKN-1 requires upstream histidine catabolism enzymes and likely proceeds through the buildup of a metabolic intermediate. (A) Schematic of *haly-1* genomic locus labelling novel alleles from EMS screen (top, arrows). A conserved active site is labelled in red and two existing alleles are labelled (bottom). Scale bar, 500 bp. (B) Fluorescent images of SKN-1 reporter animals in a wildtype or *amdh-1(uth29)* mutant background fed RNAi targeting *haly-1* and *uroc-1*. Scale bar, 100 μm. (C) Quantification of SKN-1 activation (*amdh-1* mutant normalized to median of wild type) from (B), Data shown are representative of n = 3 biological replicates with n > 250 animals per condition for each replicate. **** = P < 0.0001 using a one-way ANOVA. (D) Fluorescent images of SKN-1 reporter animals in a wildtype or *amdh-1(uth29)* mutant background with (+ His) or without (control) 10mM histidine added to the media. Scale bar, 100 μm. (E) Quantification of SKN-1 activation (*+*His normalized to median of control) from (D), Data shown are representative of n = 3 biological replicates with n > 141 animals per condition for each replicate. **** = P < 0.0001 Mann-whitney U test. (F) Fluorescent images of SKN-1 reporter animals in an *amdh-1(uth29)* mutant background fed RNAi with or without 10mM histidine added to the media. Scale bar, 100 μm. (G) Quantification of SKN-1 activation (*+*His normalized to median of control) from (F), Data shown are representative of n = 3 biological replicates with n > 193 animals per condition for each replicate. **** = P < 0.0001 one-way ANOVA.

AMDH-1 is the third enzyme in the catabolism of histidine to glutamate. Two upstream enzymes conserved in *C. elegans*, HALY-1 and Y51H4A.7, renamed to **<u>UROC</u>**anate hydratase protein <u>1</u>, UROC-1, catalyze histidine to glutamate conversion through the formation of two intermediate catabolites, urocanate and IP (fig 1b). One possible mechanism, supported by the finding that *haly-1* mutants suppress SKN-1 activation in *amdh-1* mutants, is that catabolite buildup activates SKN-1. To explore this possibility, we performed epistasis experiments using RNAi to knock down the upstream enzymes of the histidine catabolism pathways in *amdh-1* mutant animals. If a buildup of the second catabolite, IP, leads to SKN-1 activation, knockdown of *uroc-1* would also suppress SKN-1 activation. We found that RNAi knockdown of *haly-1* phenocopies the suppression seen in *haly-1* mutants and that *uroc-1* completely suppressed the activation of SKN-1 in *amdh-1* mutants (fig 3b,c). These findings support the hypothesis that SKN-1 activation in these mutants proceed through a catabolite intermediate, likely IP.

### Histidine supplementation amplifies SKN-1 activation in AMDH-1 mutants

The poor solubility and short half-life of catabolites upstream of *amdh-1*, urocanate and IP, prevented the direct testing of catabolite activation. Rather, we tested whether increasing flux through the histidine catabolism pathway differentially affects *amdh-1* mutants compared to wild type animals. Notably, this differential activation would depend on one or both of the upstream enzymes, *haly-1* and *uroc-1*, if the mechanism proceeds through a catabolite intermediate. Accordingly, we supplemented SKN-1 reporter animals, in both wild-type and *amdh-1* mutant backgrounds, with histidine. Strikingly, *amdh-1* mutant animals exhibit a robust activation of SKN-1 upon histidine supplementation when compared to wildtype animals (fig. 3d,e). Indeed, this activation is partially suppressed by knockdown of *haly-1* and completely suppressed by knockdown of *Y51H4A.7/uroc-1* (Fig 3f,g). These data further support a model that the catabolite intermediate IP drives SKN-1 activation.

### *amdh-1* mutants are sensitive to heat stress and resistant to oxidative stress

SKN-1 can be either beneficial or detrimental to organismal physiology and aging depending on expression level. For example, moderate activation of SKN-1 extends lifespan while high expression can shorten lifespan (Paek et al., 2012; Tullet et al., 2008). Moreover, activation of SKN-1 increases oxidative stress survival while decreasing survival to other stressors such as heat (Crombie et al., 2016; Deng et al., 2020). Considering the strong induction of SKN-1 upon *amdh-1* loss of function, we questioned whether this activation was beneficial or detrimental to organismal health. First, we assessed the lifespans of *amdh-1(uth29)* mutants compared to wild-type animals and found no significant effect on adult lifespan (fig. 4a). We next tested whether SKN-1 activation affects the animal’s ability to survive different stress conditions. Predictably, *amdh-1* mutants are resistant to tert-butyl hydroperoxide, an organic peroxide known to induce the OxSR, likely representing a “priming” effect that SKN-1 has on these animals to survive oxidative stress (fig. 4b). Surprisingly, we observed a stark change in resistance to thermal stress, a 50% decrease in thermotolerance compared to wildtype animals. The decreased thermotolerance of *amdh-1* mutants was completely rescued by RNAi knockdown of *skn-1*, suggesting that SKN-1 activation is detrimental to thermotolerance (fig. 4c). Additionally, stress survival on tunicamycin was also modestly reduced (fig. 4d). Notably, we did not observe a change in any other fluorescent reporters that measure the activation of other cellular stress responses (supplemental figure 4a-e). Together, our data suggest that SKN-1 activation in *amdh-1* mutants drives a physiological tradeoff that preserves survival under oxidative stress conditions at the cost of heat and ER stress resilience without a significant impact on these well defined transcriptional stress responses.

**Figure 4.**
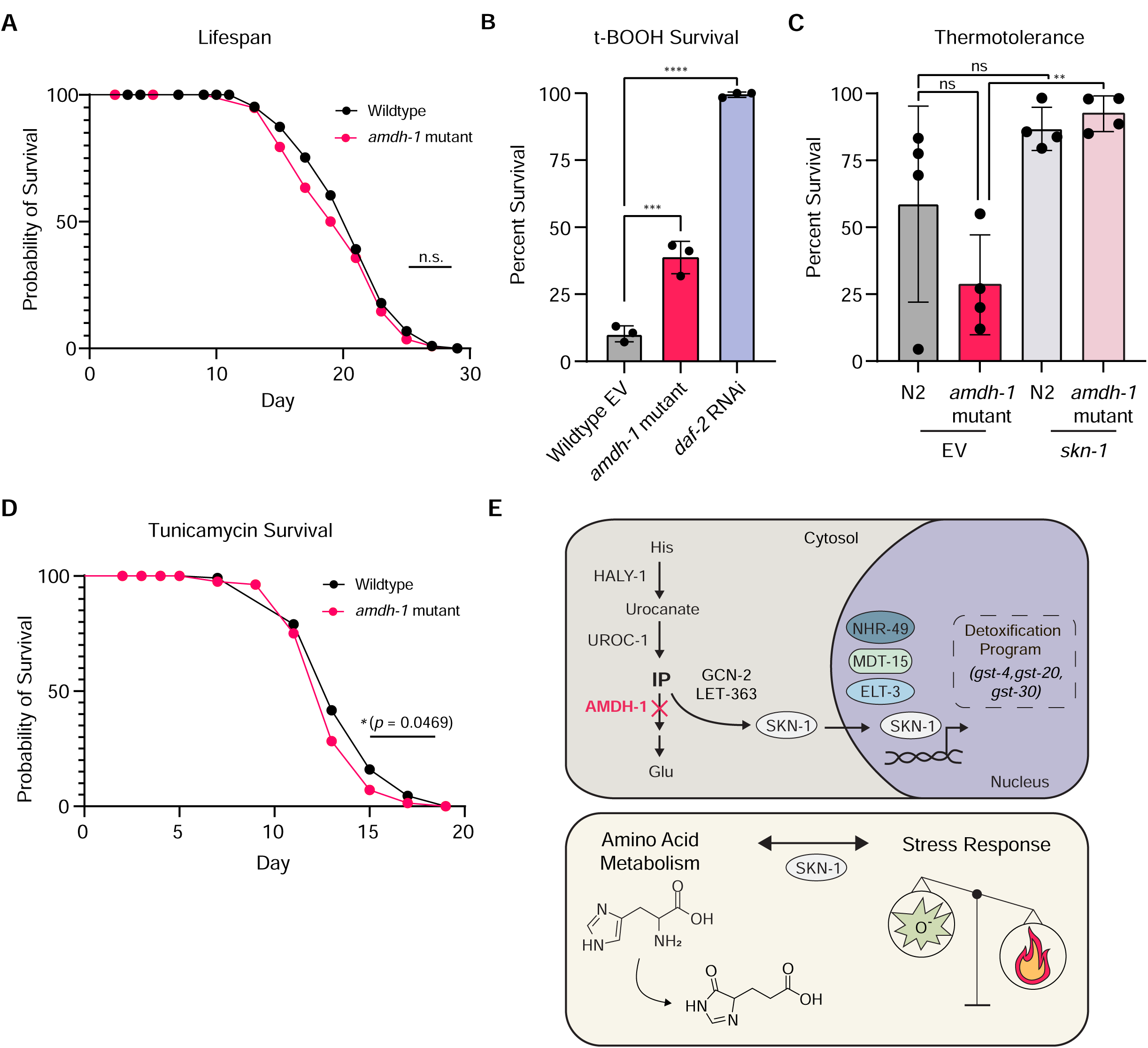
Physiological consequences of SKN-1 activation in *amdh-1* mutants. (A) Survival of wildtype animals (N2) and *amdh-1(uth29)* mutant worms at 20°C. Each data point represents one biological replicate of n > 40. One-way ANOVA with Šídák’s multiple comparisons test, ** = P < .01. (B) Survival of animals on plates containing 7.5mM t-BOOH for 16 hours. One-way ANOVA with Šídák’s multiple comparisons test, *** = P < 0.001, **** = P < 0.0001 (C) Thermotolerance of animals shifted to heat shock temperature (34°C) for 14-15 hours. One-way ANOVA with Šídák’s multiple comparisons test, ** = P < .01 (D) Survival of animals on 25ng/μL tunicamycin plates (E) Schematic of findings. *Top* - Perturbation of histidine catabolism via *amdh-1* mutation leads to a buildup of a catabolite that, directly or indirectly, activates a transcriptional program of detoxification enzymes driven by SKN-1. This response requires nuclear factors NHR-49, MDT-15 and ELT-3. *Bottom* - Activation of SKN-1 via perturbation of histidine catabolism leads to a physiological tradeoff of increased oxidative stress resistance and decreased heat tolerance

## DISCUSSION

### SKN-1 as a metabolic surveillance factor

Previous studies have reported that perturbation of tryptophan, threonine, or proline catabolism evoke a SKN-1-mediated transcriptional response (Fisher et al., 2008; Pang et al., 2014; Tang and Pang, 2016; Ravichandran et al., 2018). Our work expands this list to include glycine, valine, phenylalanine and histidine catabolism as surveillance targets of SKN-1, which can have direct effects on stress resilience. Further, we demonstrate that knockdown of a conserved amidohydrolase in the histidine catabolism pathway leads to a buildup of a catabolite, likely IP, to activate a transcriptional response driven by SKN-1. This response appears partially dependent on nutrient regulators, GCN-2 and mTORC2, and OxSR regulators ELT-3, NHR-49 and MDT-15 (fig. 4e). It is unknown whether SKN-1 activation in this context is a direct or indirect consequence of catabolite buildup, however the activation of SKN-1 results in a clear trade-off of increased oxidative stress resilience for increased sensitivity to heat stress.

Interestingly, existing literature suggests multiple amino acid catabolites can signal to SKN-1. For example, proline catabolism is modulated upon pathogen exposure to accumulate the intermediate pyrroline-5-carboxylate (P5C) to activate SKN-1 through ROS production (Tang and Pang, 2016). Other reports have demonstrated that the perturbation of tyrosine catabolism, through mutation in fumarylacetoacetate hydrolase, *fah-1*, causes stunted growth and intestinal degradation through tyrosine catabolites (Fisher et al., 2008). Intriguingly, this mutation also activates a SKN-1-dependent transcriptional reporter, and these phenotypes can be reversed through knockdown of upstream enzymes (Fisher et al., 2008). Our findings help unify these previously observed phenomena and suggest that SKN-1 is a metabolic surveillance factor that can integrate information from multiple catabolite activators into transcriptional programs to affect physiology.

Importantly, amino catabolism pathways are implicated in disease progression and cancer treatment. A previous study has shown that perturbation of this pathway decreases sensitivity to the chemotherapeutic methotrexate and increasing flux through this pathway has been proposed to increase methotrexate efficacy in patients (Kanarek et al., 2018). While simple dietary intervention is appealing, our understanding of the consequences of catabolite buildup remains incomplete. Indeed, one study indicates that histidine supplementation can cause hepatic enlargement in patients with liver disease (HoleČek, 2020). Moreover, in type I tyrosinemia, tyrosine catabolite buildup is thought to be a main source of damage to proteins and DNA, contributing to pathology (Ferguson et al., 2010). We observe that histidine catabolites can also signal to effectors, such as SKN-1, to modulate physiology. Thus, further studies to uncover the clear role of catabolite intermediates as important modulators of cell signaling is at the forefront of understanding human disease.

### 4-Imidazolone-5-Propanoate (IP) as a signaling catabolite

Enzymes of the histidine catabolism pathway are highly conserved from bacteria to humans (Bender, 2012). Interestingly, mutations in the conserved IPase in *Klebisella aerogenes* are innocuous unless supplemental histidine is added and upstream enzymes are active, in which case the bacteria are poisoned and fail to grow (Boylan and Bender, 1984). This phenotype bears striking resemblance to the phenotype observed here, in which *C. elegans* mutants for the IPase, *amdh-1*, show exaggerated phenotypes when supplemented with histidine, dependent on the upstream enzymes *haly-1* and *uroc-1*. To date, no direct experimentation has shown the toxicity of this catabolite, likely due to the short half-life of IP (Bowser Revel and Magasanik, 1958; Rao and Greenberg, 1961). IP may either directly or indirectly activate SKN-1 through a breakdown product, production of oxidative stress or through a distinct mechanism. Indeed, 4-imidazolones have previously been shown to cause oxidative stress and make up many advanced glycation end-products (AGEs), biomarkers that correlate with aging and metabolic disease (Niwa et al., 1997; Omar et al., 2018).

### Requirements of SKN-1 activation in *amdh-1* mutants

To date, the MAPK signaling pathway has been reported to control nearly all instances of SKN-1 activation in *C. elegans* (Blackwell et al., 2015). Here we report a SKN-1 transcriptional response to altered histidine catabolism that is independent of conserved MAPK components *sek-1* and *pmk-1*. Although the identity of upstream signaling components remain elusive, several kinases known to influence SKN-1 activation remain as candidates downstream of the signaling catabolite (Kell et al., 2007). Notably, we found that knockdown of *gcn-2* partially suppressed SKN-1 activation in *amdh-1* mutants. *gcn-2* is a conserved protein kinase which functions in the integrated stress response (ISR) (Pakos-Zebrucka et al., 2016). Interestingly, the homolog of SKN-1, NRF2, has known functions in regulating the transcription of ATF4, the core effector of the ISR, in mammals (Pakos-Zebrucka et al., 2016). Moreover the *gcn-2* inhibitor, *impt-1*, extends the lifespan of *C. elegans* and requires SKN-1, highlighting potential interactions between *gcn-2*, the ISR and SKN-1. Further studies are needed to determine the entirety of the signaling cascade that culminates in the activation of SKN-1 upon metabolic perturbations, as the partial dependence of *gcn-2* suggests other factors are involved. Interestingly, we identified *suco-1* as a suppressor of SKN-1 activation in *amdh-1* mutants. *suco-1* is a homolog of the SLP1 protein in yeast, which is hypothesized to participate in protecting nascent proteins from degradation during folding in the ER (Zhang et al., 2017). Previous work has identified UPR^ER^ components such as *ire-1* and *hsp-4* in the transcriptional response to oxidants arsenite and tBOOH (Glover-Cutter et al., 2013). If *suco-1* functions in a similar capacity in *C. elegans* as it does in yeast, this could implicate other ER protein homeostasis pathways in the regulation of SKN-1. Indeed, an ER-associated isoform of SKN-1, SKN-1A, is known to be a monitor of proteasome function and may modulate crosstalk between the ER and SKN-1 (Lehrbach et al., 2017)

### Activation of SKN-1 initiates a physiological trade off

Titration of SKN-1 expression is important for the pro-longevity nature of this transcription factor. Moderate overexpression of SKN-1 or mutation of the negative regulator *wdr-23* extends the lifespan of *C. elegans*, while hypomorphic mutants or worms treated with *skn-1* RNAi exhibit a shortened lifespan (Grushko et al., 2021; Ganner et al., 2019; Tullet et al., 2017; Tang and Choe, 2015; Tullet et al., 2008). However, gain-of-function animals with constitutive expression of SKN-1 and animals containing high-copy arrays of SKN-1 exhibit a mild decrease in lifespan (Paek et al., 2012; Tullet et al., 2008). Additionally, activation of SKN-1 is beneficial for oxidative stress survival while detrimental to survival under other conditions such as heat, ER or mitochondrial stress (Deng et al., 2020). SKN-1 activation upon metabolic perturbation, as shown here, provides short term benefit to survive oxidative stress but comes at the cost of sensitivity to heat and ER stress. This may represent a physiological trade off, where SKN-1 prioritizes the allocation of cellular resources to defend against a perceived threat at the cost of sensitivity to other perturbations. Identification and study of the physiologically relevant consequences of SKN-1 activation will be crucial to understanding how modulation of this master transcription factor may be leveraged to affect human disease.

## Supporting information

Table 1 - Amino Acid Catabolism Screen

Table S1 - RNAseq Data

## AUTHOR CONTRIBUTIONS

PAF designed and executed experiments and wrote the manuscript. RHS executed early experiments and intellectually contributed to project design/execution. TS and TB executed stress response reporter experiments. RBZ assisted with RNA-sequencing analysis. HKG and HZ assisted with early experiments crucial to the direction of the project. AD provided funding and substantial intellectual support. All authors assisted with the editing of the manuscript.

## ACKNOWLEDGEMENTS

We thank Larry Joe for NGS library preparation. We thank all Dillin lab members for useful discussion throughout the project and feedback on the manuscript. We also would like to thank Dr. Robert Bender (University of Michigan) for thoughtful discussions throughout the project. This work was supported by the following grants: PAF was supported by 4T32GM007232-40 through the NIH. R.H.S is supported by 4R00AG065200-03 through the NIA. R.B.Z. is supported by the Larry L. Hillblom Foundation Fellowship 2019-A-023-FEL. HKG is supported by NSF Grant # DGE175814 and NIA 1F99AG068343-01. H.Z. is supported by the Larry L. Hillblom Foundation Fellowship 2020-A-018-FEL. This work used the Vincent J. Coates Genomics Sequencing Laboratory at UC Berkeley, supported by NIH S10 OD018174 Instrumentation Grant. AD and the lab are funded by R01ES021667 and R01AG059566 from the NIH and the Howard Hughes Medical Institute. Some strains were provided by the CGC, which is funded by NIH Office of Research Infrastructure Programs (P40 OD010440).

## DECLARATION OF INTERESTS

No competing interests to declare.

## DATA AND AVAILABILITY

The raw RNA-seq data were uploaded to the NCBI short read archive (PRJNA801069). Access for reviewers is available at https://dataview.ncbi.nlm.nih.gov/object/PRJNA801069?reviewer=nchldc4t4gjac3ff5rr3k8g7rn.

## METHODS

### Strain List

| <u>Strain</u> | <u>SOURCE</u> | <u>IDENTIFIER</u> |
| --- | --- | --- |
| *C. elegans*: strain N2 (Bristol) | CGC | N2 |
| *C. elegans*: strain LD1171: *ldIs3* [*gcs-1p::GFP* + *rol-6(su1006)*] | CGC | LD1171 |
| *C. elegans*: strain CL2166: *dvIs19*[*pAG15(gst-4p::GFP::NLS)*] III | CGC | CL2166 |
| *C. elegans*: strain AM446: *rmIs223*[*pC12C8.1::GFP*; *rol-6(su1006)* II] | Gift from Morimoto lab (Morley and Morimoto, 2004) | AM446 |
| *C. elegans*: strain AGD3126: *zcIs13*[*hsp-6p::GFP*] | This paper | AGD3126 |
| *C. elegans*: strain AGD3111: *dvIs19*[*pAG15(gst-4p::GFP::NLS)*] III; *sek-1(km4)* X | This paper | AGD3111 |
| *C. elegans*: strain AGD3013: *uth93 ;amd-1(uth29)*III; | This paper | AGD3013 |
| dvIs19[pAG15(gst-4p::GFP::NLS)]; uthEx967[haly-1p::haly-1::haly-1 3' UTR, myo-3p::mCherry] |  |  |
| *C. elegans*: strain AGD2530: uth93 ;*amd-1*(uth29)III; dvIs19[pAG15(gst-4p::GFP::NLS)] | This paper | AGD2530 |
| *C. elegans*: strain AGD2529: uth92 ;*amd-1*(uth29)III; dvIs19[pAG15(gst-4p::GFP::NLS)] | This paper | AGD2529 |
| *C. elegans*: strain AGD2432: pmk-1(km25) IV; dvIs19[pAF15(gst-4p::GFP::NLS)] | This paper | AGD2432 |
| *C. elegans*: strain AGD2307:*amd-1*(uth29)III; dvIs19[pAG15(gst-4p::GFP::NLS)] | This paper | AGD2307 |
| *C. elegans*: strain AGD2306: *amd-1*(uth29)III; *rmIs223[pC12C8.1::GFP; rol-6(su1006) II]* | This paper | AGD2306 |
| *C. elegans*: strain AGD2053: *zcls4[hsp-4p::GFP]V* (SJ4005 5x backcross) | This paper | AGD2053 |
| *C. elegans*: strain AGD1848: *amd-1*(uth29)III | This paper | AGD1848 |

#### *C. elegans* maintenance

All *C. elegans* strains were maintained at 15°C on NGM plates with OP50 *E. coli* B strain. All experiments were performed at 20°C on RNAi plates (NGM agar, 1 mM IPTG, 100 μg/mL carbenicillin) with HT115 *E. coli* K12 strain bacteria containing the RNAi plasmid pL4440 empty vector as a negative control (EV) or containing sequence to synthesize a double-stranded RNA against a target gene unless otherwise stated. All RNAi constructs were isolated from the Vidal or Ahringer RNAi library and sequence verified before using. For all experiments, eggs were obtained using a standard bleaching protocol (1.8% sodium hypochlorite and 0.375 M KOH) and arrested at the L1 stage overnight in M9 (22 mM KH_2_PO_4_ monobasic, 42.3 mM Na_2_HPO_4_, 85.6mM NaCl, 1 mM MgSO_4_) without food for synchronization. The next day, synchronized L1 animals were placed on HT115 bacteria and grown until day 1 of adulthood. For histidine supplementation experiments, plates were supplemented with 10mM histidine that was buffered with HCl to pH 7.0. Animals were grown on histidine supplemented plates after L1 synchronization for the duration of the entire experiment.

*haly-1* rescue experiments were performed using a ∼4.3kb amplicon from the genomic DNA of N2 (bristol) animals to include a putative promoter region (∼1.1 kb) and complete CDS (∼2.6kb, including introns) flanked on each side by the endogenous 5’ and 3’ UTRs as annotated by wormbase (∼0.3kb each). This amplicon was generated in a standard PCR reaction using N2 genomic DNA with the forward primer oPF388 (5’ - ttgtccaataaacctttgtcc - 3’) and reverse primer oPF389 (5’ -tccatataaccctgtaactcc - 3’) and sequenced verified using standard Sanger sequencing after PCR purification. Array positive animals were generated by injecting *haly-1(uth92)* or *haly-1(uth93)* animals with purified amplicon at 40 ng/μL along with a co-injection marker (myo-3p::mCherry) at 5 ng/μL. Two independent arrays were isolated from different parent animals for each *haly-1* mutant allele.

### Lifespan and Stress Assays

Lifespan measurements were assessed on RNAi plates (standard NGM agar supplemented with 1mM IPTG and 100ug/mL carbenicillin) with HT115 bacteria carrying pL4440 empty vector RNAi. Worms were synchronized by standard bleaching/L1 arresting as described and kept at 20°C throughout the duration of the experiment. Adult worms were moved away from progeny onto fresh plates for the first 5-7 days until progeny were no longer visible and scored every 1 to 2 days until all animals were scored. Animals with bagging, vulval explosions, or other age-unrelated deaths were censored and removed from quantification. For tunicamycin survival assays, animals were moved onto tunicamycin (25 ng/μl) or 1% DMSO plates at D1 of adulthood and scored as described for standard lifespan measurements. For thermotolerance, worms were synchronized by bleaching as described above, L1 arrested, and plated on RNAi plates with HT115 bacteria carrying pL4440 empty vector or other RNAi. At D1, 15-20 worms per plate with 3-4 plates per condition were exposed to 34°C heat via incubator for 14-15 hours. Plates were then removed from the incubator and manually assessed for movement and pharyngeal pumping, using light head taps where necessary, to determine survival. Worms that displayed internal hatching (bagging) or crawled onto the side of the plate and desiccated were censored and omitted from the final analysis. Percent alive was calculated using the number of living worms divided by the total number of worms excluding censored animals for each strain. For oxidative stress survival, worms were bleach synchronized, L1 arrested, and plated on RNAi plates with HT115 bacteria carrying empty vector or *daf-2* RNAi. At D1, ∼100 animals per condition were transferred to 4-5 NGM plates containing 7.5mM t-booh (Luperox TBH70X, Sigma). Worms were scored for survival every 2 hours, starting at 12 hours, until the 16 hour time point.

### Fluorescence imaging and quantification

Image acquisition was performed as previously described (Bar-Ziv et al., 2020). Briefly, day 1 animals were picked under a standard dissection microscope onto a solid NGM plate that contained a ∼15μL drop of 100 nM sodium azide. Immobilized worms were aligned head to tail and images were captured on an Echo Revolve R4 microscope equipped with an Olympus 4x Plan Fluorite NA 0.13 objective lens, a standard Olympus FITC filter (ex 470/40; em 525/50; DM 560), and an iPad Pro for the camera and to drive the ECHO software.

To quantify fluorescence, a COPAS large particle biosorter was used as previously described (Bar-Ziv et al., 2020). Data were collected gating for size (time of flight [TOF] and extinction) to exclude eggs and most L1 animals. Data were processed by censoring events that reached the maximum peak height for Green or Extinction measurements (PH Green, PH Ext = 65532) and censoring events < 300 TOF to exclude any remaining L1 animals. For the reporters with low basal fluorescence (AGD2053, AGD3126), data > 0 were included. For reporter strains with visible basal fluorescence (CL2166, AGD2307), data >= 10 were included for subsequent statistical analysis. All fluorescence data were normalized to TOF to account for worm size. For all *amdh-1(uth29)* mutant experiments, ‘SKN-1 activation’ was quantified by normalizing Green/TOF value for each mutant to the median of the wildtype population for each condition. For the MAPK mutant experiments (fig. 2a), ‘SKN-1 activation’ was quantified by normalizing the Green/TOF value for each mutant fed *amdh-1* RNAi to the median of that mutant fed EV RNAi. All data is plotted as log(FC) SKN-1 activation.

### RNA isolation, sequencing and analysis

Animals were bleach synchronized and grown to Day1 adulthood on empty vector RNAi plates. At least 2,000 animals per condition per replicate were washed off plates using M9 and collected. After a 30 second spin down at 1,000 RCF, M9 was aspirated, replaced with 1mL Trizol, and the tube was immediately frozen in liquid nitrogen to be stored at -80°C for downstream processing. RNA was harvested after 3 freeze thaw cycles in liquid nitrogen/37°C water bath. After the final thaw, 200μL (1:5 chloroform:Trizol) of chloroform were added to the sample, vortexed, and the aqueous phase was collected after centrifugation in a gel phase lock tube. RNA was isolated from the obtained aqueous phase using a Qiagen RNeasy MiniKit according to manufacturer’s directions. Library preparation was performed using Kapa Biosystems mRNA Hyper Prep Kit (Roche, product number KK8581) using dual index adapters (KAPA, product number KK8722). Sequencing was performed using Illumina HS4000, mode SR100, through the Vincent J. Coates Genomic Sequencing Core at University of California, Berkeley.

For RNA-seq analysis of *amdh-1(uth29)* mutants, the raw sequencing data were uploaded to the Galaxy project web platform and the public server at usegalaxy.org was used to analyze the data (Afgan et al., 2016). Paired end reads were aligned using the Kallisto quant tool (Version 0.46.0) with WBcel235 as the reference genome. Fold changes and statistics were generated using the DESeq2 tool with Kallisto quant count files as the input. Volcano plots were generated using the GraphPad Prism version 8.0.0 for Windows (GraphPad Software, San Diego, California USA, www.graphpad.com) on the fold change and adjusted-p values generated by the DESeq2 analysis (table s1). For analysis of previously published data, raw reads were downloaded from the Gene Expression Omnibus (GEO), (Accession: GSE123531 and GSE63075) and analyzed as described above.

### EMS mutagenesis screen to find suppressors of SKN-1 activation

*amdh-1(uth29); gst-4p::GFP::NLS* (strain AGD2307) were mutagenized to find suppressors of *gst-4p::GFP* signal. Briefly, ∼150 L4 animals were picked into a 1.5mL eppendorf tube containing 1mL M9 buffer and spun down at 1,000 RPM for 1 minute. The M9 was aspirated from the tube and replaced with 1mL fresh M9 and spun again. To the washed worm pellet, 5μL of EMS was added, the tube was parafilmed and left nutating at 20C for 4 hours. After incubation, the worms were spun down and rinsed 4 times with 1mL M9. Waste was collected and neutralized with 1:1 KOH before discarding. Rinsed, mutagenized, worms were plated overnight. The next day, mutagenized worms were picked onto 10 large plates seeded with OP50 bacteria, 10 per plate, and allowed to lay eggs for 24 hours. Three days later, adult F1 animals were bleached and plated onto fresh large plates. F2 mutants were screened for suppression of *gst-4p::GFP* under a fluorescent microscope compared to age matched, un-mutagenized, conrol animals.

Genomic DNA was extracted from mutants of interest using the Puregene Cell and Tissue Kit (Qiagen), as previously described (Lehrbach et al., 2017). 2ug of purified DNA was sheared using a Covaris S220 focused-ultrasonicator to produce ∼400 bp fragments. Library preparation was performed with 1ug of sheared DNA using Kapa Biosystems Hyper Prep Kit (Roche, product number KK8504) dual index adapters (KAPA, product number KK8727). Sequencing was performed using the Illumina NovaSeq6000 platform through the Vincent J. Coates Genomic Sequencing Core at University of California, Berkeley. Raw reads were uploaded to the Galaxy project web platform and the public server at usegalaxy.org was used to analyze the data (Afgan et al., 2016). Reads were aligned using the Bowtie2 tool with WBcel235/ce11 as the reference genome. The MiModD tool suite (Baumeister lab) was used on the Variant Allele Contrast (VAC) mapping mode to call, extract and filter variants to compare mutants to the parental, un-mutagenized strain. Causative genes were identified through a combination of genetic complementation, deep sequencing and RNAi phenocopy experiments.

## Notes

https://dataview.ncbi.nlm.nih.gov/object/PRJNA801069?reviewer=nchldc4t4gjac3ff5rr3k8g7rn.

### Competing Interest Statement

The authors have declared no competing interest.

