## Supplemental Figures and Legends for "SKN-1 is a metabolic surveillance factor that monitors amino acid catabolism to control stress resistance"

#### Supplemental Figure 1

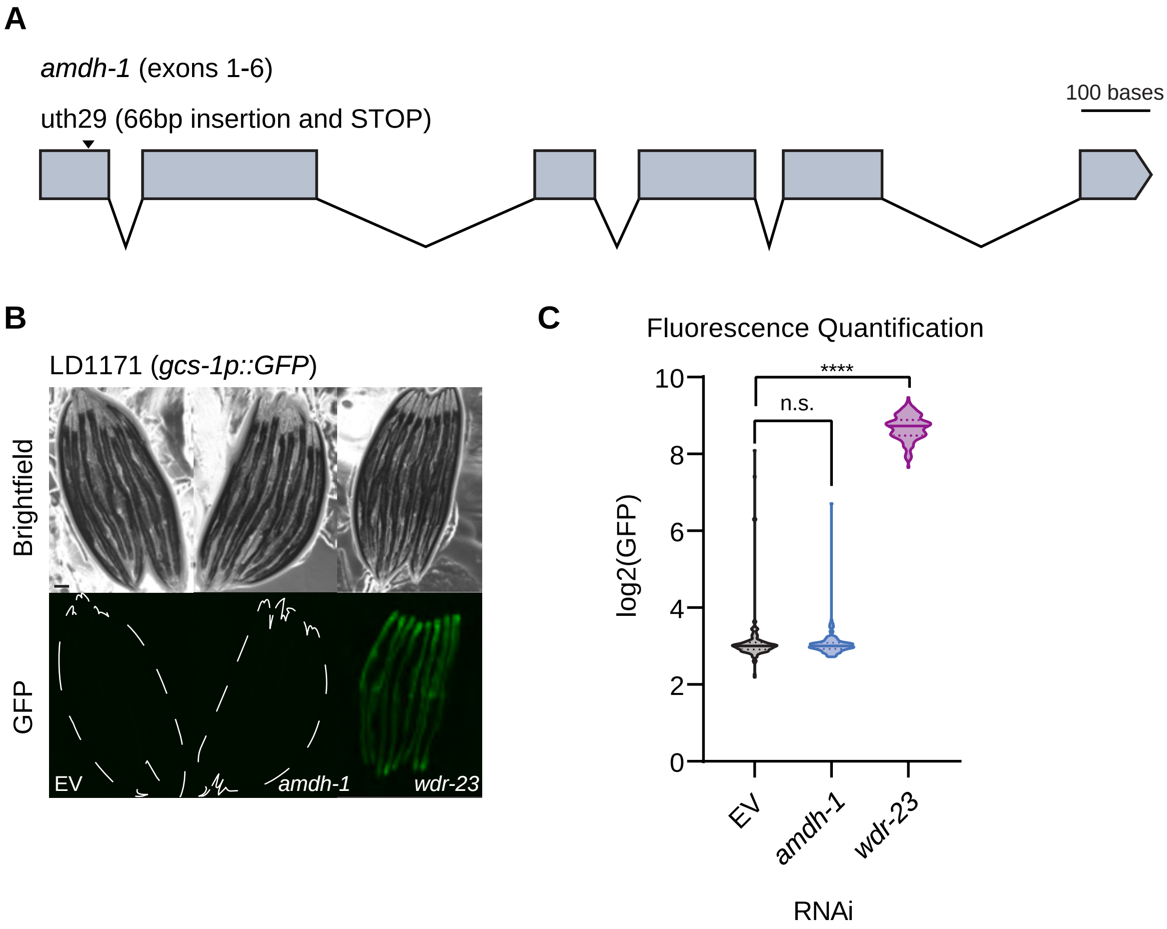

### Supplemental Figure 2

**A**

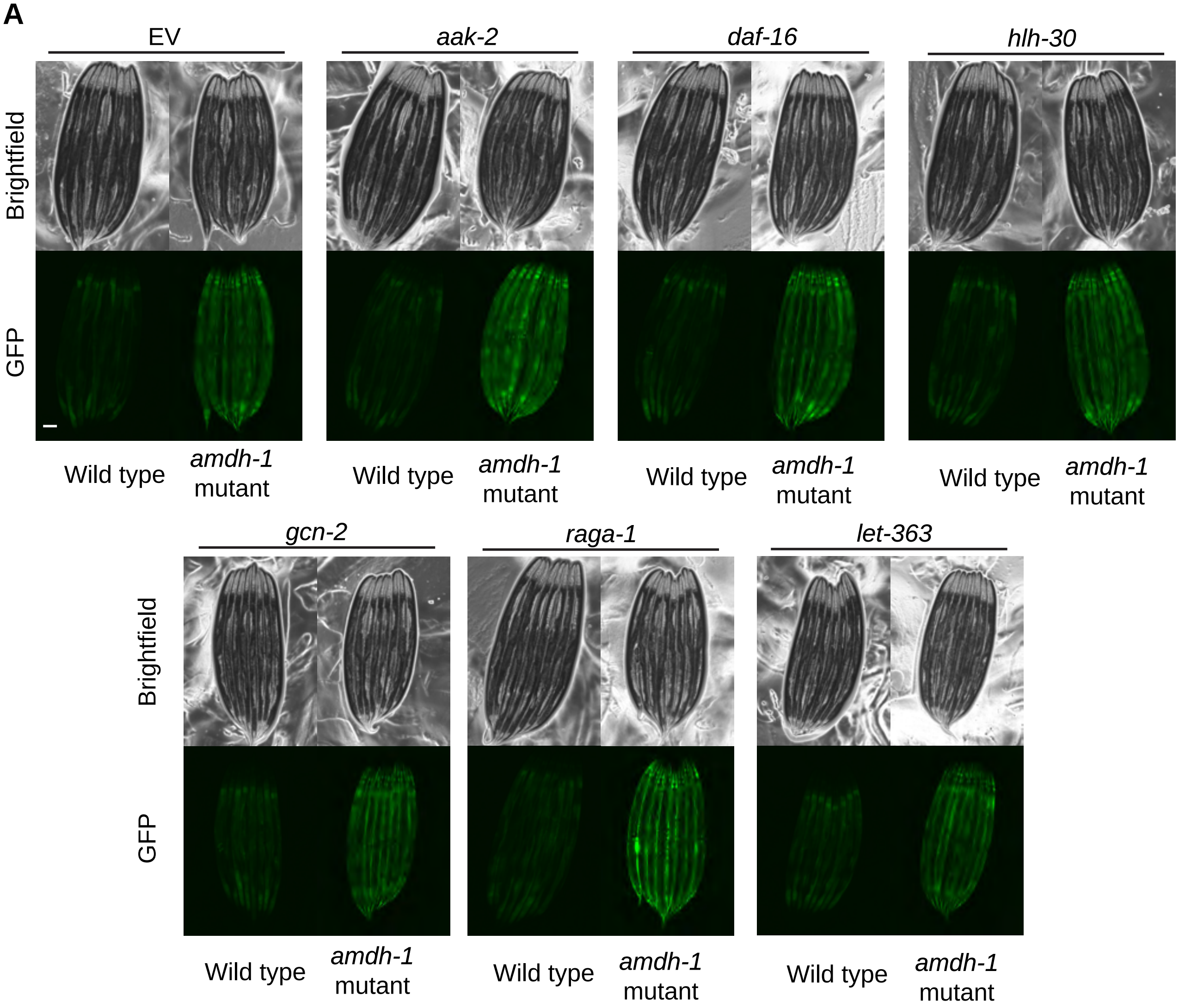


**B**

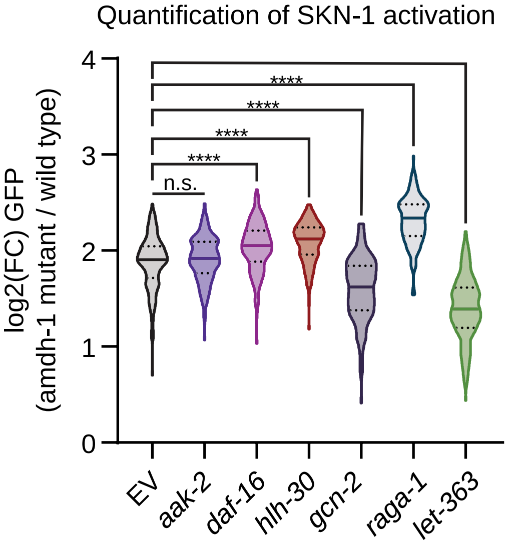


**C**

| **Allele** | **Gene** | **Mutation** |
|---|---|---|
| *uth89* | *skn-1* | A514T |
| *uth94* | *elt-3* | Q128* |
| *uth112* | *suco-1* | A429M |
| *uth92* | *haly-1* | G164E |
| *uth93* | *haly-1* | G265E |
| *uth95* | *haly-1* | G551V |

**Figure 4 supplemental**

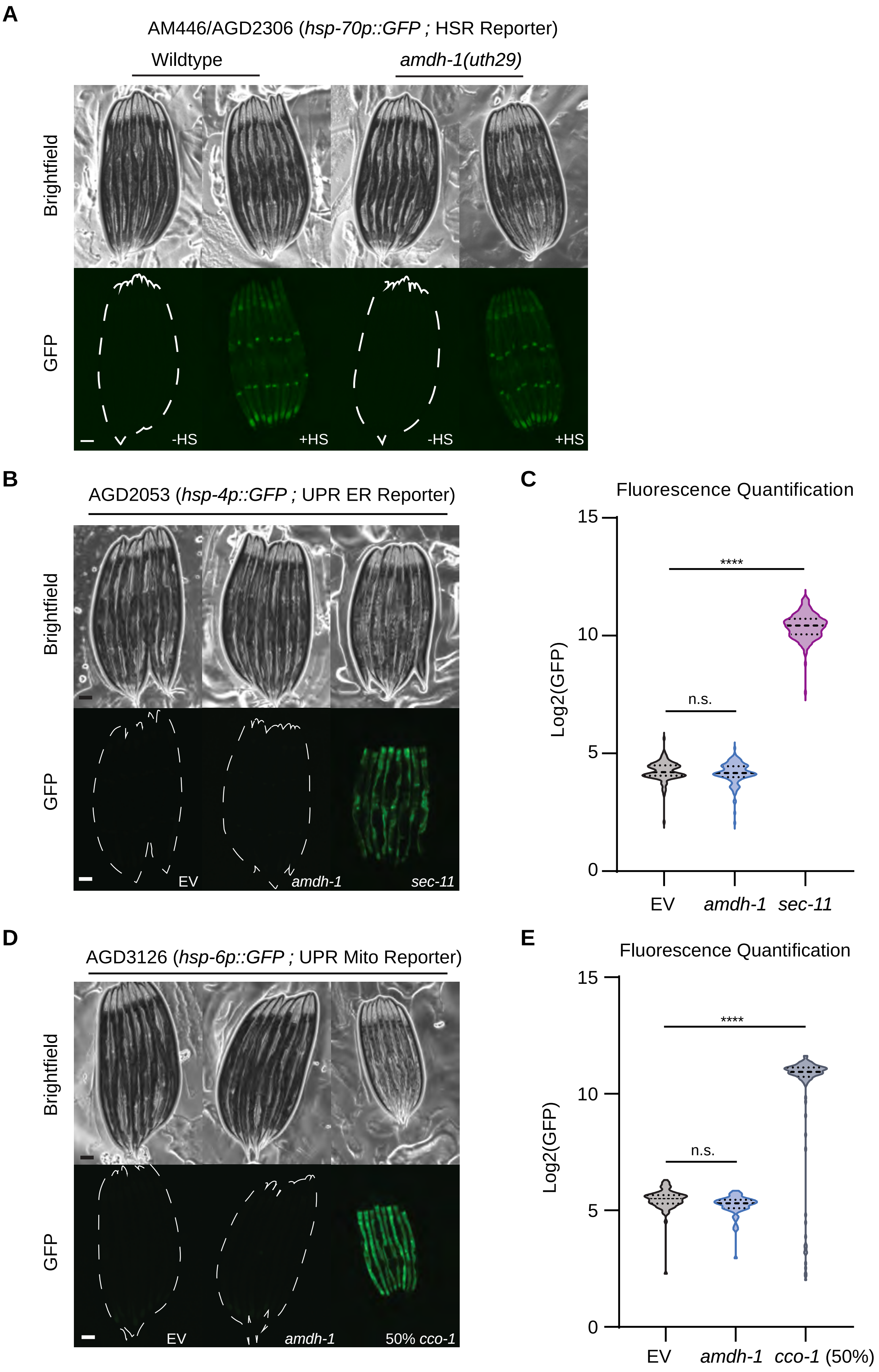

**Supplemental Figure 1**

(A) Schematic of *amdh-1/T12A2.1* genomic locus showing the location of allele *uth29* engineered using CRISPR/Cas9 (top, arrow). Scale bar, 100 bases. (B) Fluorescent images of SKN-1 reporter worms (*gcs-1p::GFP)* fed RNAi targeting *amdh-1* and *wdr-23*. Scale bar, 100 µm. (C) Quantification of (B), Data shown are representative of $n = 4$ biological replicates with $n > 60$ animals per condition for each replicate. **** = $P < 0.0001$, n.s. = not significant using a one-way ANOVA non-parametric test (Kruskal-wallis).

**Supplemental Figure 2**

(A) Fluorescent images of SKN-1 reporter animals (*gst-4p::GFP)* fed RNAi targeting. Scale bar, 100 µm. (B) Quantification of (A), Data shown are representative of $n = 2$ biological replicates with $n > 123$ animals per condition for each replicate. (C) Table of causative mutations mapped using whole genome sequencing.

**Supplemental Figure 3**

(A) Schematic of *haly-1* genomic DNA (gDNA) rescue array used for genetic rescue experiments to confirm causality (B) Fluorescent images of *haly-1; amdh-1* double mutants with or without *haly-1* gDNA rescue array. Scale bar, 100 µm.

#### Supplemental Figure 3

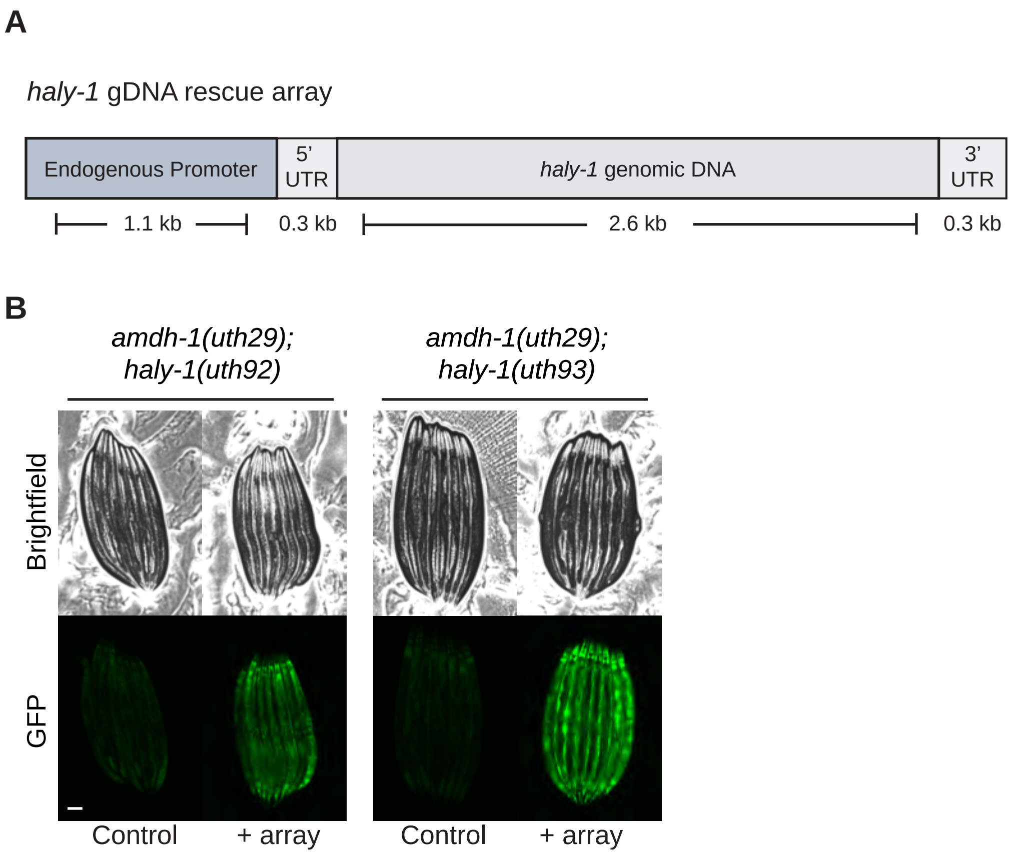

**Supplemental Figure 4**

(A) Fluorescent images of heat shock response (HSR) reporter animals (hsp-70p::GFP) with or without heat shock treatments (34C) in wildtype and amdh-1(uth29) mutant animals. Data shown are representative of n = 3 biological replicates. Scale bar, 100 µm. (B) Fluorescent images of a reporter of the unfolded protein response of the ER (UPR ER) animals (*hsp-4p::GFP)* fed RNAi. Scale bar, 100 µm. (C) Quantification of (B), Data shown are representative of n = 3 biological replicates with n > 50 animals per condition for each replicate. **** = P < 0.0001, n.s. = not significant using a one-way ANOVA non-parametric test (Kruskal-wallis) (D) Fluorescent images of UPR Mito reporter animals (*hsp-6p::GFP)* fed RNAi. Scale bar, 100 µm. (E) Quantification of (D), Data shown are representative of n = 3 biological replicates with n > 50 animals per condition for each replicate. **** = P < 0.0001, n.s. = not significant using a one-way ANOVA non-parametric test (Kruskal-wallis)
